# Replication of MERS and SARS coronaviruses in bat cells offers insights to their ancestral origins

**DOI:** 10.1101/326538

**Authors:** Susanna K. P. Lau, Rachel Y. Y. Fan, Hayes K. H. Luk, Longchao Zhu, Joshua Fung, Kenneth S. M. Li, Emily Y.M. Wong, Syed Shakeel Ahmed, Jasper F. W. Chan, Raven K. H. Kok, Kwok-Hung Chan, Ulrich Wernery, Kwok-Yung Yuen, Patrick C. Y. Woo

## Abstract

Previous findings of Middle East Respiratory Syndrome coronavirus (MERS-CoV)-related viruses in bats, and the ability of *Tylonycteris*-BatCoV HKU4 spike protein to utilize MERS-CoV receptor, human dipeptidyl peptidase 4 hDPP4, suggest a bat ancestral origin of MERS-CoV. We developed 12 primary bat cell lines from seven bat species, including *Tylonycteris pachypus*, *Pipistrellus abramus* and *Rhinolophus sinicus* (hosts of *Tylonycteris*-BatCoV HKU4, *Pipistrellus*-BatCoV HKU5 and SARS-related-CoV respectively), and tested their susceptibilities to MERS-CoVs, SARS-CoV and human coronavirus 229E (HCoV-229E). Five cell lines, including *P. abramus* and *R. sinicus* but not *T. pachypus* cells, were susceptible to human MERS-CoV EMC/2012. However, three tested camel MERS-CoV strains showed different infectivities, with only two strains capable of infecting three and one cell lines respectively. SARS-CoV can only replicate in *R. sinicus* cells, while HCoV-229E cannot replicate in any bat cells. Bat dipeptidyl peptidase 4 (DPP4) sequences were closely related to those of human and non-human primates but distinct from dromedary DPP4 sequence. Critical residues for binding to MERS-CoV spike protein were mostly conserved in bat DPP4. DPP4 was expressed in the five bat cells susceptible to MERS-CoV, with significantly higher mRNA expression levels than those in non-susceptible cells (*P*=0.0174), supporting that DPP4 expression is critical for MERS-CoV infection in bats. However, overexpression of *T. pachypus* DPP4 failed to confer MERS-CoV susceptibility in *T. pachypus* cells, suggesting other cellular factors in determining viral replication. The broad cellular tropism of MERS-CoV should prompt further exploration of host diversity of related viruses to identify its ancestral origin.

## IMPORTANCE

Existing evidence suggests that MERS-CoV may be originated from bats. In particular, *Tylonycteris*-BatCoV HKU4 spike protein was shown to utilize MERS-CoV receptor, human dipeptidyl peptidase 4 (DPP4). To better understand the potential infectivities of MERS-CoV in bats, we developed 12 primary bat cell lines from seven bat species, including *Tylonycteris pachypus* and *Rhinolophus sinicus* (host of *Tylonycteris*-BatCoV HKU4 and SARS-related-CoV respectively). MERS-CoV demonstrated much broader cellular tropism than SARS-CoV and HCoV-229E, being able to infect five cell lines, including *R. sinicus* but not *T. pachypus* cells. The close phylogenetic relationship between bat and human DPP4 genes supported the ability of MERS-CoV to infect bat cells, while DPP4 expression appeared critical for MERS-CoV replication. However, overexpression of *T. pachypus* DPP4 failed to confer MERS-CoV susceptibility in *T. pachypus* cells. The broad cellular tropism of MERS-CoV may reflect the host diversity of MERS-CoV-related viruses and offers insights into its ancestral origin.

## INTRODUCTION

Coronaviruses (CoVs) are important pathogens in animals and humans, responsible for a variety of respiratory, enteric, hepatic and neurological diseases. They are now classified into four genera, *Alphacoronavirus*, *Betacoronavirus*, *Gammacoronavirus* and *Deltacoronavirus*, with *Betacoronavirus* further divided into lineages A to D (1–6). Humans are infected by six CoVs, including *human CoV 229E* (HCoV-229E) and *human CoV NL63* (HCoV-NL63) belonging to *Alphacoronavirus*; human CoV OC43 (HCoV-OC43) and *human CoV HKU1* (HCoV HKU1) belonging to *Betacoronavirus* lineage A; *Severe Acute Respiratory Syndrome-related CoV* (SARSr-CoV) belonging to *Betacoronavirus* lineage B; and *Middle East Respiratory Syndrome CoV* (MERS-CoV) belonging to *Betacoronavirus* lineage C (7–16). The emergence potential of CoVs is believed to be related to their tendency for mutation and recombination, allowing the generation of new viruses being able to adapt to new hosts (5, 17–24).

Bats are an important reservoir of alphacoronaviruses and betacoronaviruses which may jump to other animals or humans to cause new epidemics (4, 25). For example, SARS-CoV is likely a recombinant virus originated from horseshoe bats as the primary reservoir and palm civet as the intermediate host (22, 26–33) Since the SARS epidemic, numerous other novel CoVs from humans or animals have been discovered (4, 34–40), allowing a better understanding of the evolutionary origin of emerging CoVs.

Although dromedary camels are now known to be the immediate animal source of the recent MERS epidemic, the evolutionary origin of MERS-CoV remains obscure (41–43) When the virus was first discovered, it was found to be closely related to *Tylonycteris* bat CoV HKU4 (Ty-BatCoV HKU4) and *Pipistrellus* bat CoV HKU5 (Pi-BatCoV HKU5) previously discovered in lesser bamboo bat (*Tylonycteris pachypus*) and Japanese pipistrelle (*Pipistrellus abramus*) respectively in Hong Kong (5, 14, 15, 44, 45). Five other lineage C betacoronaviruses closely related to MERS-CoV were subsequently detected in bats, including BtVs-BetaCoV/SC2013, Hp-BatCoV HKU25 from China bats and Coronavirus Neoromicia/PML-PHE1/RSA/2011 (NeoCoV), BtCoVNeo5038/KZN/RSA/2015 and BatCoV PREDICT/PDF-2180 from African bats (46–50). Besides bats, a lineage C betacoronavirus, Erinaceus CoV VMC/DEU, subsequently defined as a novel species, Hedgehog coronavirus 1, has also been discovered in European hedgehogs, a group of animals being phylogenetically closely related to bats (51). While none of these animal viruses represents the immediate ancestor of MERS-CoV, the spike protein of Ty-BatCoV HKU4 was most closely related to that of MERS-CoV, and was shown to utilize the MERS-CoV receptor, human dipeptidyl peptidase 4 (hDPP4) or CD26, for cell entry (52, 53). Previous studies have also shown that MERS-CoV was able to infect bat cell lines and Jamaican fruit bats (54, 55). These findings suggest that bats may be the primary host of the ancestor of MERS-CoV.

Although MERS-CoV has been shown to replicate in various animal cell lines including bat cells (54–59), the broad tissue tropism was mainly demonstrated using type strain EMC/2012, and no comparison was made between different MERS-CoV strains. Moreover, cells from the bat hosts of MERS-CoV-related viruses, such as *T. pachypus* and *P. abramus* which harbor Ty-BatCoV HKU4 and Pi-BatCoV HKU5 respectively, were not included in previous studies, which may be due to the geographical limitation of these bat species. To better understand the replicative ability of MERS-CoV in bat cells, which may provide clues on the origin of MERS-CoV, we developed diverse primary bat cell lines from different bat species, including *Rhinolophus sinicus* (the host of SARSr-BatCoV) and *T. pachypus* (the host of Ty-BatCoV HKU4), and tested their susceptibilities to infection by different strains of MERS-CoV, SARS-CoV and HCoV-229E. The DPP4 mRNA sequences of six bat species and their expression in bat cells were determined to correlate with viral replication results. Our findings showed differential cell tropism between different strains of MERS-CoV, SARS-CoV and HCoV-229E, which offers insights to the origin of MERS-CoV.

## RESULTS

### Five of 12 tested bat cell lines are susceptible to MERS-CoV EMC/2012 infection

Since lineage C betacoronaviruses closely related to MERS-CoV were detected in bats, we developed 12 diverse primary bat cell lines from seven different bat species, including *Hipposideros pomona*, *Miniopterus pusillus*, *Myotis ricketti*, *Pipistrellus abramus* (the host of Pi-BatCoV HKU5), *Rhinolophus sinicus* (the host of SARSr-BatCoV and Rs-BatCoV HKU2), *Tylonycteris pachypus* (the host of Ty-BatCoV HKU4), *Rousettus leschenaultii* (the host of many viruses including Ro-BatCoV HKU9), which were subject to infection with MERS-CoV at multiplicity of infection (MOI) of 1. Viral titers were determined by RT-qPCR on day 5 p.i․. Five of the 12 cell lines (*M. ricketti* lung, *P. abramus* kidney, *R. sinicus* kidney and lung, and *R. leschenaulti* kidney cells) propagated MERS-CoV with at least one log_10_ increase in viral load. The highest increase in viral load was observed in *R. sinicus* kidney and lung cells, which was comparable to that observed in Vero cells (Fig. 1). Cytopathic effects (CPEs) were observed in infected *M. ricketti* lung and *R sinicus* lung cells (Fig. 2). The infectivities of the viruses from culture supernatants were confirmed by passage in Vero cells with CPE. *H. pomona* kidney, *M. pusillus* kidney, *M. ricketti* kidney, *P. abramus* lung, *R. leschenaulti* lung and *T. pachypus* kidney and lung cells did not support MERS-CoV infection.

**FIG 1.**
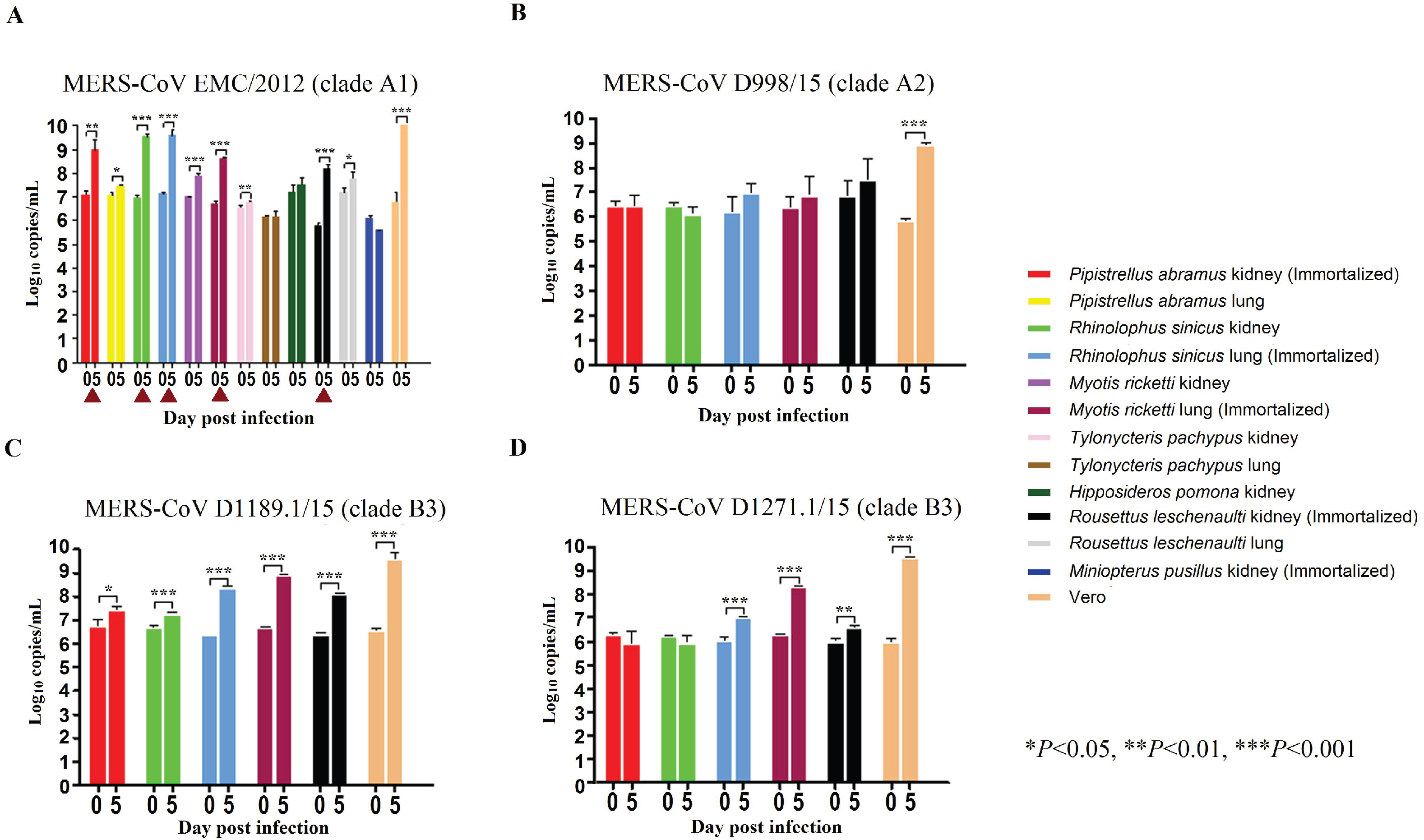
The 12 bat cell lines (PAK: *Pipistrellus abramus* kidney, PAL *Pipistrellus abramus* lung, RSK: *Rhinolophus sinicus* kidney, RSL: *Rhinolophus sinicus* lung, MRK: *Myotis ricketti* kidney, MRL: *Myotis ricketti* lung, TPK: *Tylonycteris pachypus* kidney, TPL: *Tylonycteris pachypus* lung, HPK: *Hipposideros pomona* kidney, RLK: *Rousettus leschenaultii* kidney, RLL: *Rousettus leschenaultii* lung, MPK: *Miniopteruspusillus* kidney) and Vero cells were subject to infection by MERS-CoV EMC/2012 (belonging to clade A1) with MOI of 1 (A). Culture supernatants were harvested at day 0 and 5 post infection. Viral titers were determined by real-time quantitative RT-PCR. Viral load was expressed as log10 copies/mL. Error bars indicate the standard deviation of triplicate samples. The five bat cell lines susceptible to MERS-CoV EMC/2012 infection with ≥1 log_10_ increase in viral load at day 5 were marked with red triangles. They were subject to infection by three other MERS-CoV strains isolated from camels in Dubai, D998/15 (belonging to clade A2) (B), D1189.1/15 (C) and D1271.1/15 (D) (belonging to clade B3) (B). Different MERS-CoV strains displayed different infectivities in these five bat cells. (\**P* < 0.05; \*\**P* < 0.01; \*\*\**P*<0.001)

**FIG 2.**
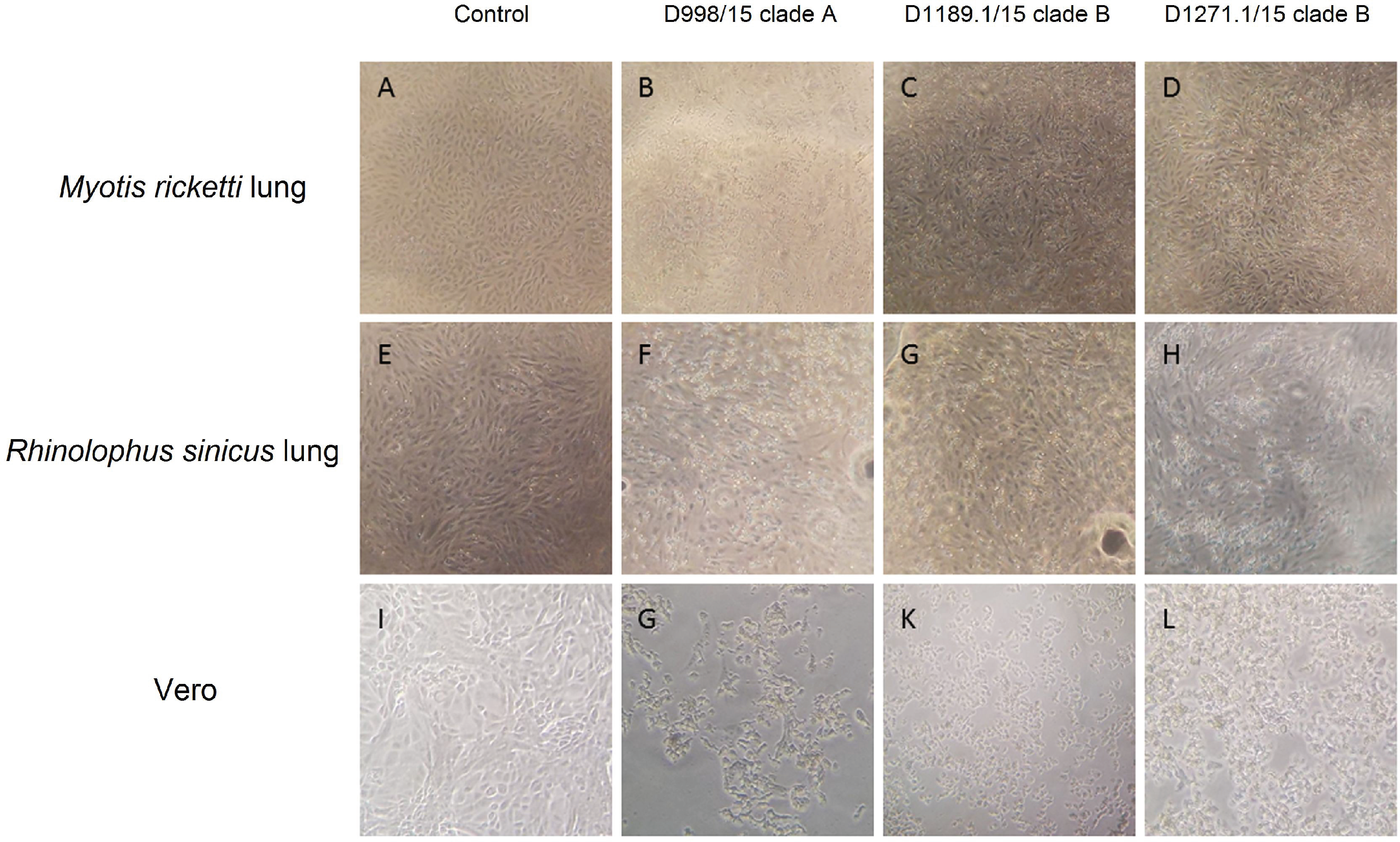
Cytopathic effects (CPE) were observed in infected *M. ricketti* lung, *R. sinicus* lung and Vero cells on 5 days post infection. CPE was compared between *Myotis ricketti* lung (immortalized) cells that were uninfected (control) (A), and infected with Dubai camel MERS strains D998/15 (B), D1189.1/15 (C) & D1271.1/15. CPE was compared between *Rhinolophus sinicus* lung (immortalized) cells that were uninfected (control) (E), and infected with Dubai camel MERS strains D998/15 (F), D1189.1/15 (G) & D1271.1/15 (H). CPE was compared between Vero cells that were uninfected (control) (I), and infected with Dubai camel MERS strains D998/15 (G), D1189.1/15 (K) & D1271.1/15 (L).

### Different MERS-CoV strains displayed different infectivities on bat cells

MERS-CoVs are currently classified into three major clades, clade A, B and C, which were further divided into subclades A1-A2, B1-B6, C1 and non-C1 (60–63). To test if different MERS-CoV strains may show similar infectivities on bat cells, the five bat cells (*M. ricketti* lung, *P. abramus* kidney, *R. sinicus* kidney and lung, and *R. leschenaulti* kidney cells), which were susceptible to MERS-CoV EMC/2012 (belonging to clade A1), were subject to infection by three other MERS-CoV strains isolated from camels in Dubai, D998/15 (belonging to clade A2), and D1189.1/15 and D1271.1/15 (both belonging to clade B3)(62). Among the five challenged bat cells, *M. ricketti* lung supported infection by both D1189.1/15 and D1271.1/15; while *R. sinicus* lung and *R leschenaulti* kidney supported infection by D1189.1/15, with at least one log_10_ increase in viral load. None of the cells supported infection by D998/15 (Fig. 1).

### SARS-CoV can replicate in *R. sinicus* cells

Since Chinese horseshoe bats are the major reservoir of SARS-related-CoVs, we also tested the 12 bat cell lines, including *R. sinicus* cells, for susceptibility to a clinical strain of SARS-CoV. SARS-CoV strain HKU-39849 can replicate in *R. sinicus* kidney but not lung cells, with at least one log_10_ increase in viral load (Fig. 3A). The other bat cells did not support SARS-CoV infection.

**FIG 3.**
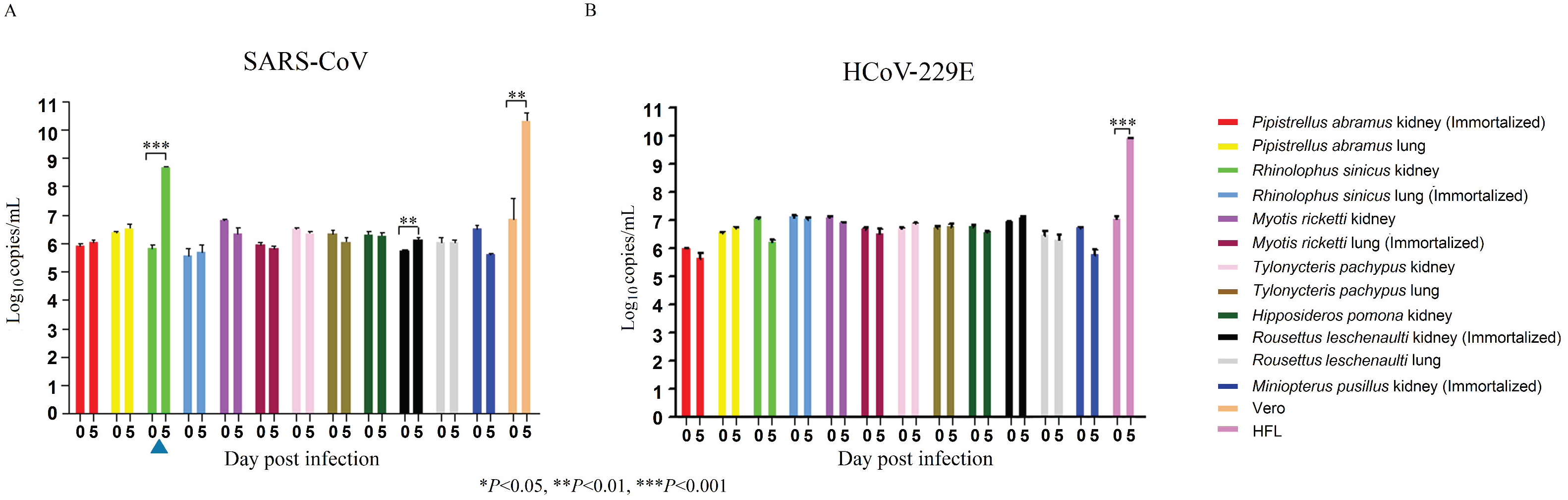
The 12 bat cell lines (PAK: *Pipistrellus abramus* kidney, PAL *Pipistrellus abramus* lung, RSK: *Rhinolophus sinicus* kidney, RSL: *Rhinolophus sinicus* lung, MRK: *Myotis ricketti* kidney, MRL: *Myotis ricketti* lung, TPK: *Tylonycteris pachypus* kidney, TPL: *Tylonycteris pachypus* lung, HPK: *Hipposideros pomona* kidney, RLK: *Rousettus leschenaultii* kidney, RLL: *Rousettus leschenaultii* lung, MPK: *Miniopterus pusillus* kidney) and Vero/HFL cells were subject to infection by SARS-CoV with MOI of 1 (A) and HCoV-229E with MOI of 0.01(B). Culture supernatants were harvested at day 0 and 5 post infection. Viral titers were determined by realtime quantitative RT-PCR. Viral load was expressed as log_10_ copies/mL. Error bars indicate the standard deviation of triplicate samples. Only RSK cells can support SARS-CoV infection with ≥1 log_10_ increase in viral load at day 5 (blue triangle) and none of the 12 bat cell lines support HCoV-229E infection. (\**P*<0.05; \*\**P*<0.01; \*\*\**P* <0.001)

### HCoV-229E cannot replicate in bat cells

The 12 bat cells were also tested for susceptible to HCoV-229E infection. HCoV-229E strain ATCC VR-740, previously isolated from a man with upper respiratory illness, cannot replicate in any of the tested bat cells (Fig. 3B).

### mRNA transcript sequence analysis and expression of DPP4 in bat cells

Partial DPP4 mRNA transcript sequences (corresponding to nt 688–1040 of hDPP4 which includes residue 229–346 where the critical residues for MERS-CoV spike protein binding are present) were determined for six of the seven bat species from which the 12 tested bat cells were developed. The sequence of *M. pusillus* was not determined, as RT-PCR for the DPP4 mRNA from bat cells was negative. Phylogenetic analysis showed that the bat DPP4 mRNA sequences formed a distinct cluster being closely related to sequences from human and non-human primates; while the dromedary camel DPP4 sequence was most closely related to that of wild boar (Fig. 4A). Previous studies have identified 14 critical residues in hDPP4 for binding of MERS-CoV spike protein (64, 65). Upon multiple sequence alignment of the corresponding regions that contain these critical residues, most of the residues are conserved in the six bat DPP4 sequences in this study (Fig. 3B). Notably, *T. pachypus* DPP4 (Tp-DPP4) contains an I→K substitution at position 295 compared to hDPP4. On the other hand, residue R336 of hDPP4 was only conserved in *T. pachypus* and *H. pomona*, both not susceptible to MERS-CoV infection.

**FIG 4.**
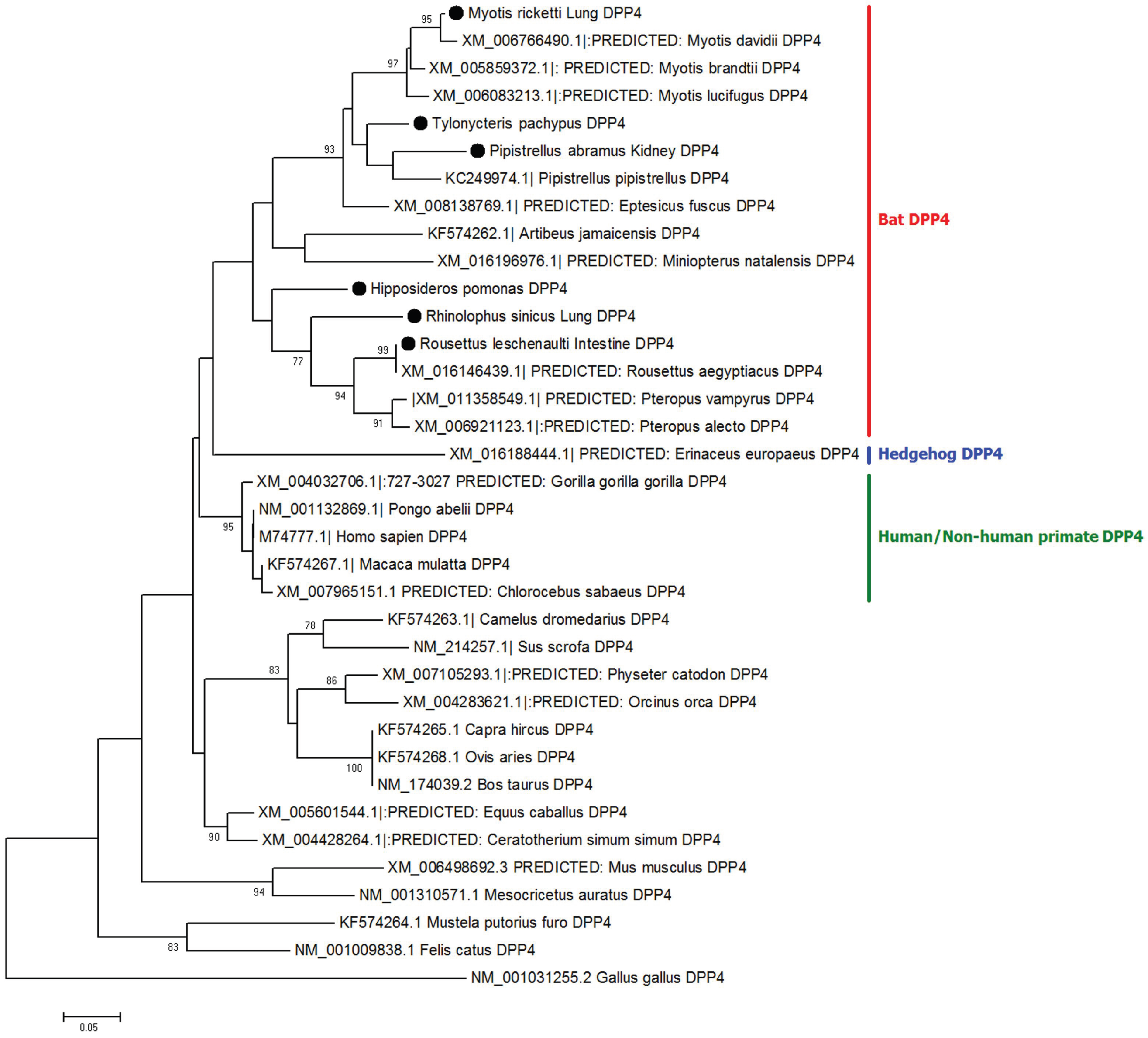
Phylogenetic analyses of partial dpp4 mRNA sequences of human, camels, bats and other animals (A). The trees were constructed by Neighbor-Joining method using JTT substitution models and bootstrap values calculated from 1000 trees. Only bootstrap values >70% are shown. 112 aa positions were included in the analyses. The scale bars represent 20 substitutions per site. Bat DPP4s that are sequenced in this study are labelled with black circles. Comparison of critical amino acid residues in DPP4 from different animal host for receptor binding in the region of residues 229–346 with respect to human DPP4 (B).

RT-qPCR of bat DPP4 mRNA in the corresponding bat cells was performed to determine the mRNA expression levels. Results showed that DPP4 mRNA was expressed in all the five bat cells that were susceptible to MERS-CoV infection, while it was also expressed in *H. pomona* kidney and *R. leschenaulti* lung cells which were not susceptible to MERS-CoV infection. The mRNA expression level in *P. abramus* kidney (susceptible to MERS-CoV) was significantly higher than that in its lung (no-susceptible) cells (*P*=0.0185 by student’s t test). Similarly, the mRNA expression level in *R. leschenaulti* kidney (susceptible to MERS-CoV) was significantly higher than that in its lung (no-susceptible) cells (*P*=0.0009 by student’s t test). Compared to bat cells that are non-susceptible, bat cells that are susceptible to MERS-CoV showed a significantly higher mRNA expression level (*P*=0.0174 by student’s t test) (Fig. 5).

**FIG 5.**
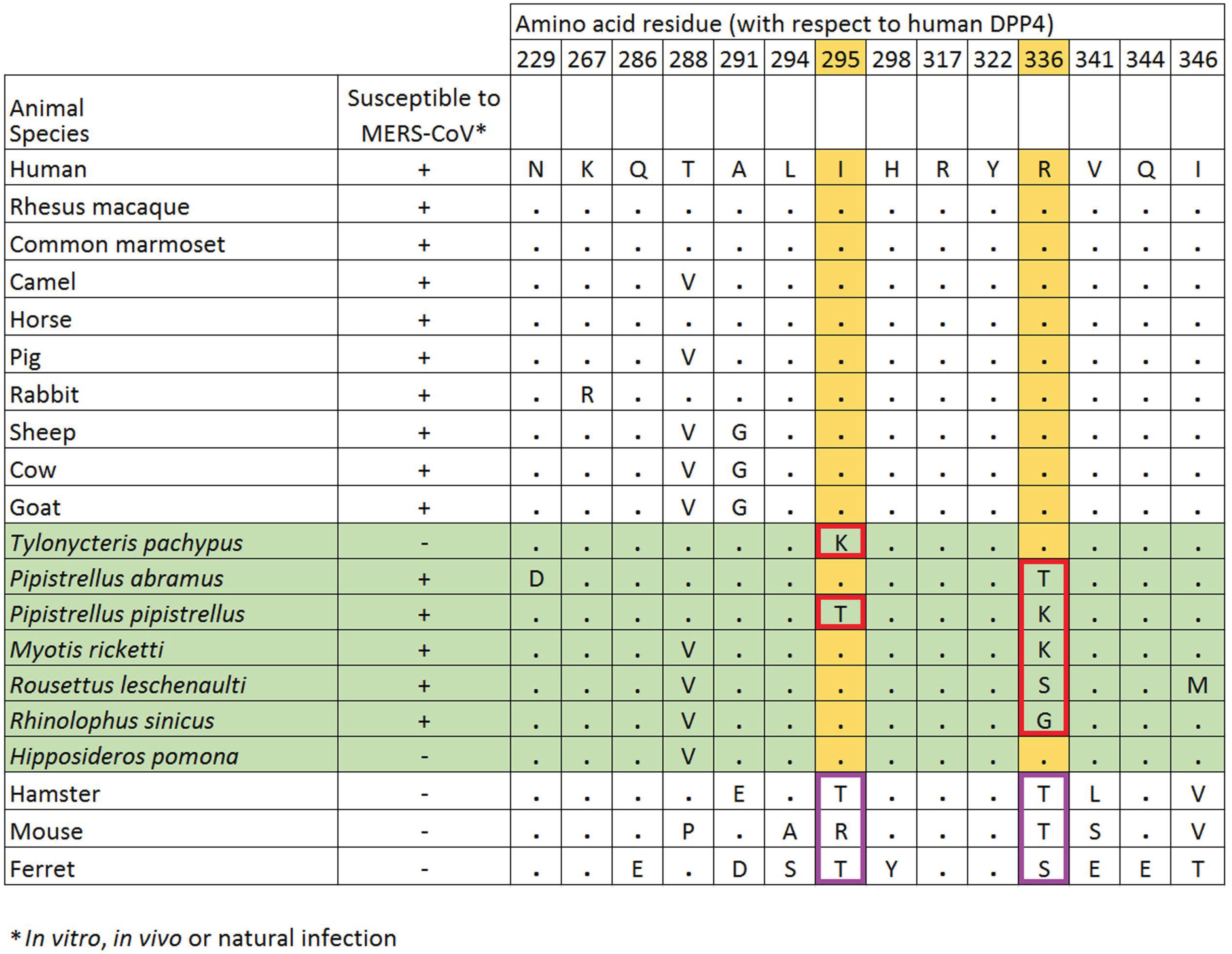
DPP4 expression analysis of bat cell lines (A). mRNA levels of DPP4 were measured in various bat cells extracts by RT-qPCR and plotted relative to Vero cells, normalized by β-actin mRNA levels. The mRNA levels of DPP4 were compared between susceptible and non-susceptible bat cell lines (B). Statistical significance was assessed by Student’s t test *P* < 0.05.

### Over-expression of Tp-DPP4 does not confer MERS-CoV EMC/2012 susceptibility to *T. pachypus* cells

Since DPP4 was not expressed in *T. pachypus* cells while this bat species hosts Ty-BatCoV HKU4, a close relative of MERS-CoV which can utilize hDPP4 for cell entry (52, 53), we overexpressed Tp-DPP4 in *T. pachypus* lung and kidney cells for infection by MERS-CoV. While a slight increase (less than one log_10_) in viral replication was noted in both mock and Tp-DPP4-overexpressed cells on day 5 p.i., no significant difference was noted between mock and Tp-DPP4-overexpressed cells (Fig. 6), suggesting that over-expression of Tp-DPP4 does not confer MERS-CoV susceptibility to *T. pachypus* cells.

**FIG 6.**
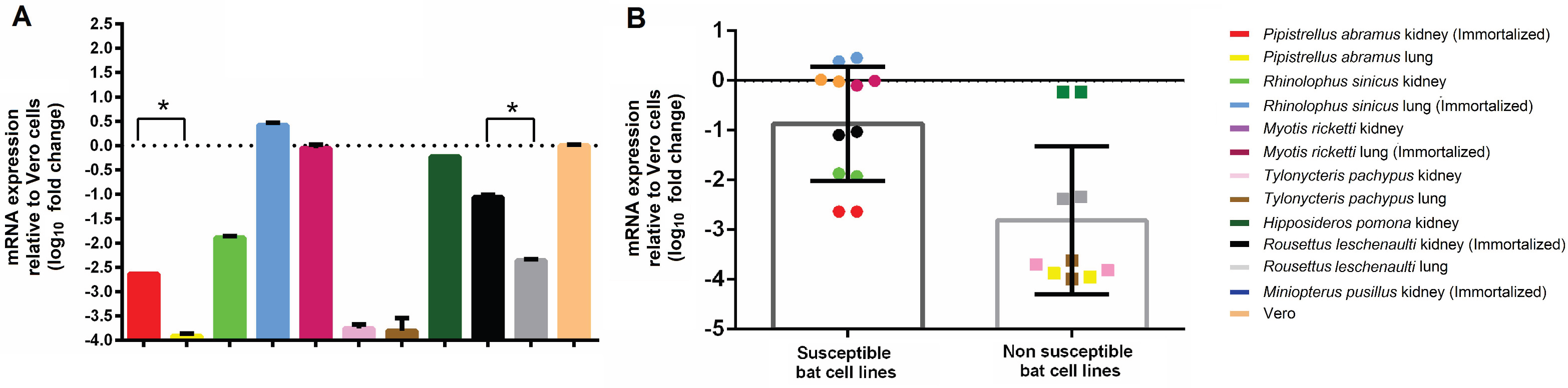
Infection assay of *T. pachypus* lung and kidney cells with tpDPP4 overexpression. Cells were infected with MERS-CoV at a multiplicity of infection (MOI) of 1 for 5 days. Determination of MERS-CoV viral load in supernatant (n = 3) by RT-qPCR with normalisation to beta-actin (represented by bar). Determination of tpDPP4 expression in cell lysates (n = 3) by RT-qPCR with normalisation to beta-actin (represented by dot). (\**P*<0.05, \*\**P*<0.01, \*\*\**P*<0.001)

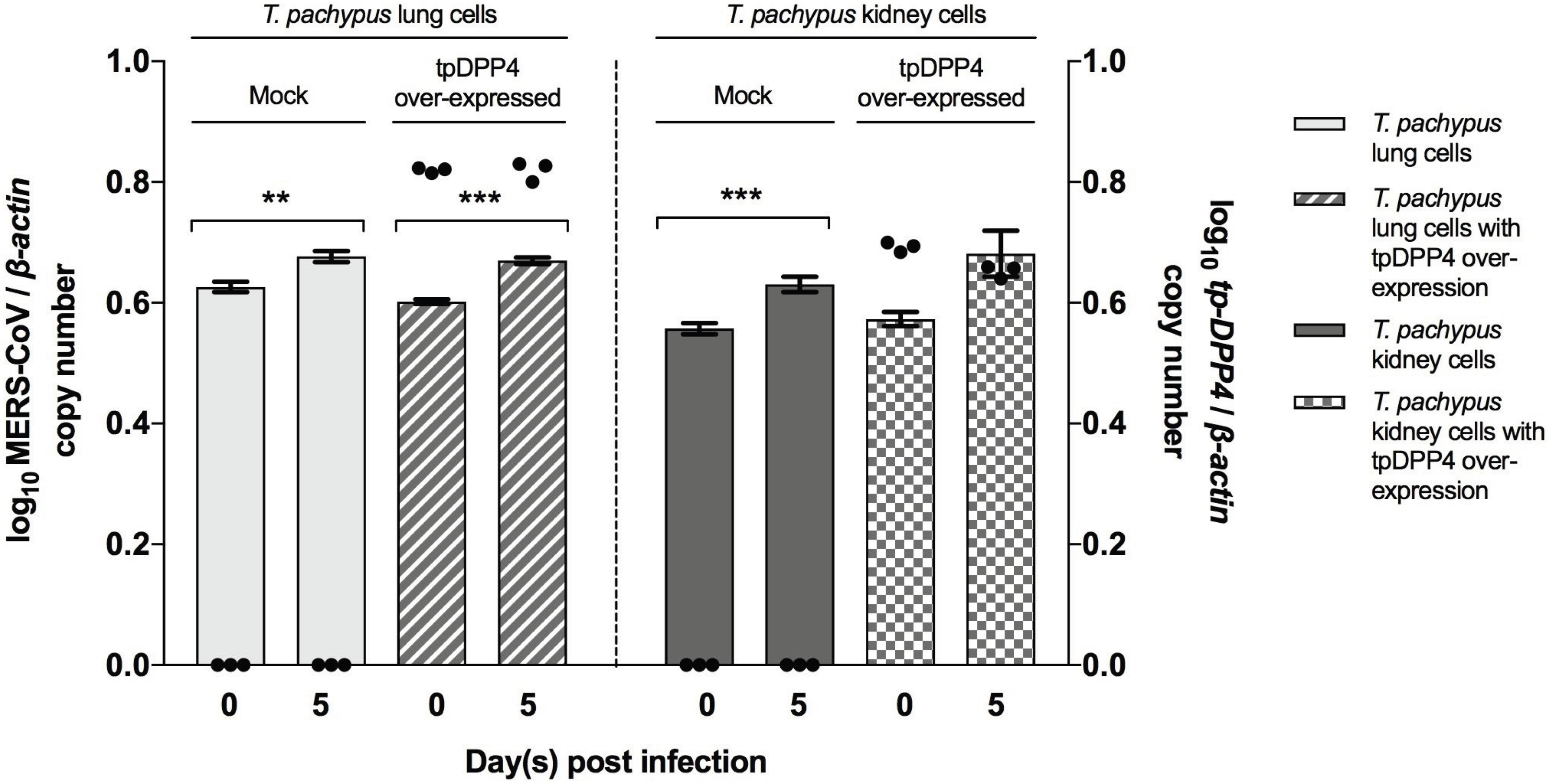

## DISCUSSION

The present study provides further support to the bat origin of MERS-CoV. Previous studies have shown that MERS-CoV can infect cell lines from different animals including cells from various bat species (54–58). However, bat cells from host species of MERS-Co-V-related viruses, such as *T. pachypus* and *P. abramus* which harbor Ty-BatCoV HKU4 and Pi-BatCoV HKU5 respectively, were not included in these studies. In this study, MERS-CoV EMC/2012 was able to replicate in cells from four different bat genera/species belonging to three different bat families including *Pteropodidae*, *Rhinolophidae* and *Vespertilionidae*. None of these bat cells were previously reported to support MERS-CoV replication. While lineage C betacoronaviruses have not been detected in bats outside the *Vespertionionidae* family, such diverse cellular tropism should prompt further studies to explore the host diversity of this group of CoVs and hence the possible evolutionary origin of MERS-CoV. In particular, the ability of MERS-CoV to replicate in cells from *P. abramus*, the host of Pi-BatCoV HKU5, may suggest that the ancestor of MERS-CoV could have originated from *Pipistrellus*-related bats. Interestingly, MERS-CoV showed the highest replicative ability in cells from *R. sinicus.* This suggests that MERS-CoV may potentially infect this bat species which is also the natural reservoir of SARSr-CoVs and animal origin of SARS-CoV (27, 29). Nevertheless, three other MERS-CoV strains belonging to either clade A2 or B3 showed differential replicative abilities in the five bat cell lines that were susceptible to EMC/2012 strain (clade A1), with strain D998/15 (clade A2) unable to replicate in all five tested cell lines. This may suggest that different MERS-CoV strains could possess different cellular tropism and host range.

The broader cellular tropism of MERS-CoV than SARS-CoV and HCoV-229E in bat cells may reflect the host diversity of lineage C betacoronaviruses in bats. Bats are now known to be the recent origin of at least two human CoVs including SARS-CoV and HCoV-229E. SARS-CoV is likely a recombinant virus arising from horseshoe bats before it jumped to civet as the intermediate host and then human (22, 31, 32, 66). SARSr-CoVs in bats were shown to utilize the SARS-CoV receptor, human angiotensin-converting enzyme 2 (hACE2), for cell entry (67). Similar to SARS-CoV, HCoV-229E is likely to have originated from bats. CoVs closely related to HCoV-229E have been detected in bats of the genus *Hipposideros* in Africa (68). More recently, CoVs even closer to HCoV-229E were identified in dromedary camels in the Middle East, which were able to utilize the receptor of HCoV-229E, human aminopeptidase N (hAPN) for cell entry (69). This suggests that dromedary camels may have served as intermediate hosts for bat-to-human transmission of HCoV-229E, which may be analogous to the evolution of MERS-CoV. Recently, a CoV, closely related to HCoV-NL63 in most genome regions except the spike protein, was detected from a bat of the genus *Triaenops* in Kenya (70). However, further studies are required to identify even closer relatives of HCoV-NL63 to ascertain its possible bat origin. In contrast to SARSr-CoVs and HCoV-229E which were mainly found in horseshoe and roundleaf bats respectively, a more diverse host range was observed in lineage C betacoronaviruses. Yet, the different bat species harboring lineage C betacoronaviruses all belonged to the family *Vespertilionidae.* Specifically, Ty-BatCoV HKU4, Pi-BatCoV HKU5, Hp-BatCoV HKU25 and BtVs-BetaCoV/SC2013 were detected in bats belonging to the genera, *Tylonycteris*, *Pipistrellus*, *Hypsugo* and *Verspetilio*, respectively in China, whereas NeoCoV and BatCoV PREDICT/PDF-2180 were detected in bats belonging to *Neoromicia* and *Pipistrellus* in Africa (47, 48). In this study, SARS-CoV could only replicate in *R. sinicus* kidney cells but not other bat cells, which reflects its evolutionary origin from horseshoe bats. On the other hand, HCoV-229E was unable to replicate in any tested bat cells including cells from *Hipposideros*. This may suggest a relatively narrow host range of SARS-CoV and HCoV-229E in bats compared to lineage C betacoronaviruses including MERS-CoV.

The close phylogenetic relationship between bat and primate DPP4 sequences may reflect the replicative ability of MERS-CoV in bat cells. Moreover, cells that supported MERS-CoV replication showed significantly higher DPP4 mRNA expression than those that were non-susceptible to MERS-CoV, suggesting that cellular DPP4 expression is critical for viral infection. Since Ty-BatCoV HKU4 from *T. pachypus* was previously shown to be able to utilize hDPP4 for cell entry (52, 53), we expected MERS-CoV to be able to utilize Tp-DPP4 for receptor binding and infect *T. pachypus* cells. While the inability of MERS-CoV to replicate in *T. pachypus* cells may be partly explained by the lack of DPP4 expression, over-expression of Tp-DPP4 was unable to confer viral susceptibility. Further studies are required to study the receptor-binding interphase between MERS-CoV and Tp-DPP4 and whether other cellular factors may play a role in determining viral replicative abilities in *T. pachypus* cells, which may offer further insights into the evolutionary origin and mechanisms of interspecies transmission of MERS-CoV. Interestingly, one of the critical residues for binding to MERS-CoV spike protein, R336, was only conserved in *T. pachypus* and *H. pomona* DPP4 (cells from these two species did not support MERS-CoV replication), but not in the other four sequenced bat DPP4 (cells from these four species supported MERS-CoV replication). This suggests that this residue may be less important for receptor binding. Binding and mutagenesis studies may help better understand the role of receptor-binding-interphase during viral evolution and interspecies jumping.

## MATERIALS AND METHODS

### Ethics statement

The collection of bat samples for developing bat cell lines was approved by the Committee on the Use of Live Animals for Teaching and Research, the University of Hong Kong, Hong Kong, China.

### Cell lines

The bat cells, Vero (African green monkey kidney) cells and HFL (human embryonic lung fibroblast) cells used in this study are described in Table 1. For development of primary bat cell lines, bats were captured in Hong Kong and euthanized before dissection for recovery of cells from organs aseptically. Briefly, kidney and lung tissue were rinsed with cold PBS and cut into small pieces. Cold 0.25% trypsin-EDTA was added to the tissues and incubated at 4°C overnight. Tissues were then incubated at 37°C on shaking platform for 30 min. Supernatants were filtered through cell strainers to remove large pieces of tissues (71) Bat cells were harvested by spinning down the supernatant at 1200 rpm for 8 min and were grown in DMEM/F12 supplemented with 15% FBS. Vero and HFL cells were grown in MEM supplemented with 10% FBS. All cells were incubated at 37°C with 5% CO_2_.

**Table 1.**
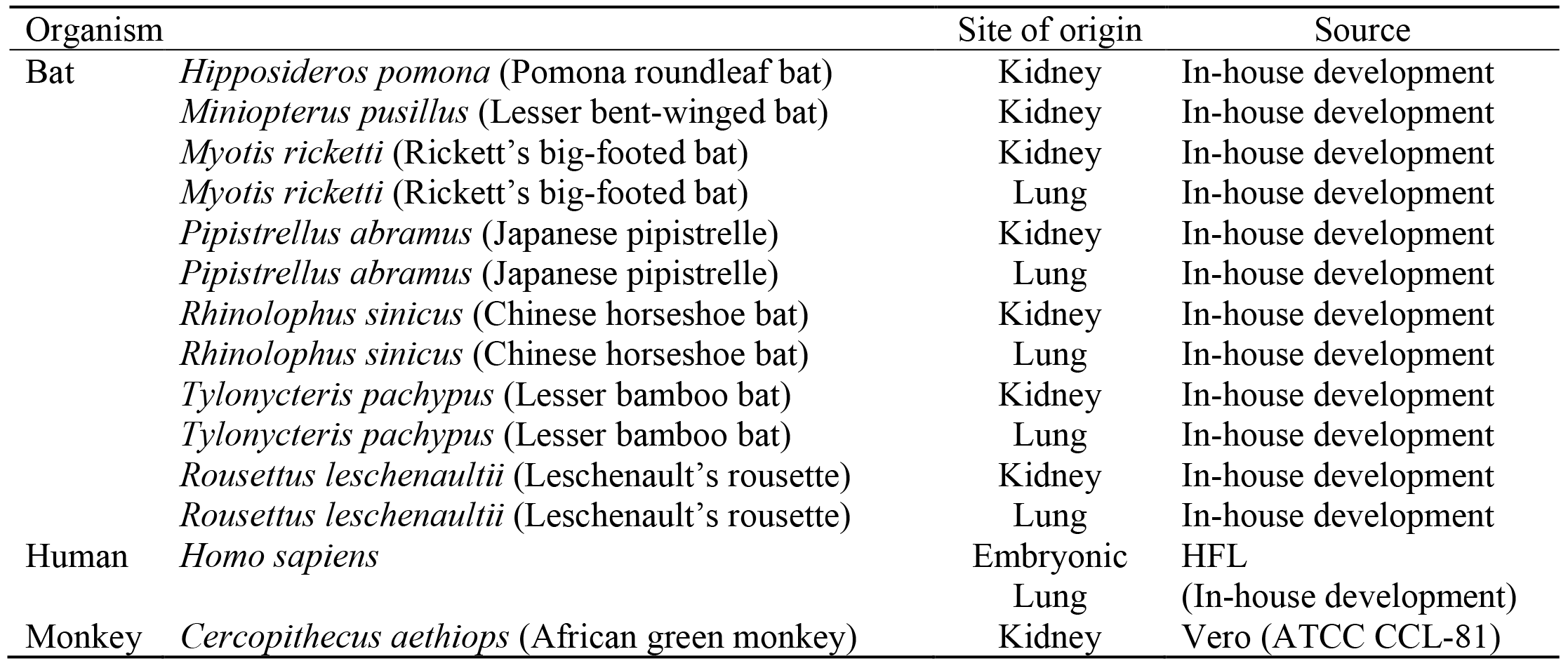
Cell lines used in the present study

### Virus isolates

MERS-CoV strain EMC/2012 was provided by Fouchier and Zaki et al (14). MERS-CoV strains D998/15, D1189.1/15 and D1271.1/15 were isolated from dromedary camels in Dubai(62). SARS-CoV strain HKU-39849 was isolated from the brother-in-law of the index patient in Hong Kong during the SARS epidemic (10). The HCoV-229E strain ATCC VR-740 was used. MERS-CoV isolates and SARS-CoV were propagated in Vero cells at MOI of 0.01 in MEM supplemented with 1% FBS. HCoV-229E was propagated in HFL.

### Infection of bat cell lines

Viral titers were determined as median tissue culture infective dose (TCID_50_) per ml in confluent cells in 96-well microtiter plates. Cells were seeded onto 24-well tissue culture plates, at 2×10^5^ cells per well with the respective medium and incubated at 37°C and 5% CO_2_ for 24 h prior to experiment. Cells were washed once with PBS and inoculated with 1 MOI of MERS-CoV or SARS-CoV, or 0.01 MOI of 229E for 1 h. After 1 h of viral adsorption, the supernatant was removed and cells were washed twice with PBS. The cells were maintained in MEM supplemented with 1% FBS for Vero and HFL cells and DMEM/F12 supplemented with 1% FBS for bat cells, before further incubation for 5 days.

### Viral replication studies

To study viral replication efficiency, progeny viruses from cell culture supernatants collected at 5 days post-infection (p.i.) were subject to reverse transcription-quantitative PCR (RT-qPCR) according to our previous protocols (72). Briefly, total RNA extracted from cell culture supernatants with QIAsymphony DSP Virus/Pathogen Mini Kit (Qiagen, Hilden, Germany) was reverse transcribed and amplified with MERS-CoV primers (forward primer 5’ -CAAAACCTTCCCTAAGAAGGAAAAG -3’; reverse primer 5’- GCTCCTTTGGAGGTTCAGACAT -3’), SARS-CoV primers (forward 5’- ACCAGAATGGAGGACGCAATG-3’; reverse 5’ -GCTGTGAACCAAGACGCAGTATTAT- 3′) and HCoV-229E primers (forward 5’-CAGTCAAATGGGCTGATGCA-3’; reverse 5’- AAAGGGCTATAAAGAGAATAAGGTATTCT-3’) using real-time one-step quantitative RT-PCR assay as described previously with modifications (56, 72). Probes for MERS-CoV [5’- (FAM)ACAAAAGGCACCAAAAGAAGAATCAACAGACC(BHQ1)-3’], SARS-CoV [5’- (FAM)ACCCCAAGGTTTACCC(NFQ)-3’] and HCoV-229E [5’- (FAM)CCCTGACGACCACGTTGTGGTTCA(BHQ1)-3’] were used (Table 2). Reactions were first incubated at 50°C for 30 min, followed by 95°C for 2 min, and were then thermal cycled for 50 cycles (95°C for 15 s, 55°C for 30 s). A series of 10 log_10_ dilutions equivalent to 1 × 10^1^ to 1 × 10^10^ copies per reaction mixture were prepared to generate calibration curves and were run in parallel with the test samples. All experiments were performed in triplicate.

**Table 2.**
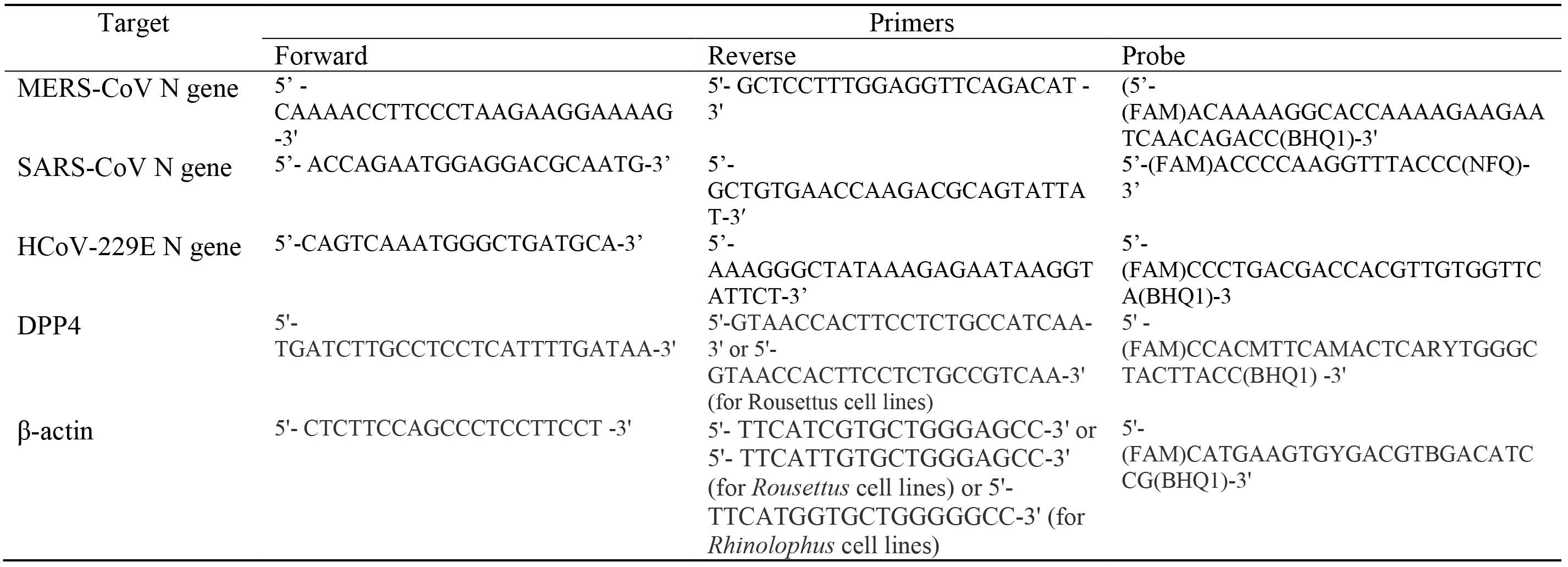
Primers used for RT-qPCR in this study

### Amplification and sequencing of partial bat DPP4 mRNA transcripts

Total RNA was extracted from bat cell lysates using RNeasy Mini Spin Column (QIAgen). cDNA was PCR amplified with primers, 5’- GTCACCAGAGGGTCATAAA-3’ and 5’- CCACTTCCTCTGCCATCAAA-3’. The PCR mixture (25 μl) contained cDNA, PCR buffer, 200 μM (each) dNTPs, and 1.0 U Iproof Polymerase (Bio-Rad, Hercules, USA). The mixtures were amplified for 40 cycles of 98°C for 10 sec, 55°C for 30 sec and 72°C for 72 sec and a final extension at 72°C for 10 min in an automated thermal cycler (Applied Biosystems, Foster City, USA). RT-PCR products were gel purified using QIAquick gel extraction kit (Qiagen), and sequenced with an ABI Prism 3700 DNA Analyzer (Applied Biosystems). The sequences obtained were compared with sequences of DPP4 genes in GenBank database. Phylogenetic tree construction was performed based on an amino acid alignment of partial DPP4 sequences (corresponding to residue 229–346 of hDPP4) using Neighbor-Joining method with JTT model by MEGA 6.0, with bootstrap values calculated from 1000 trees.

### DPP4 expression analysis

To study DPP4 expression profiles in different bat cell lines, cell lysates were collected for total RNA extraction using RNeasy Mini Spin Column (QIAgen). RNA was eluted in 50 μl of RNase-free water and was used as template for one step RT-qPCR with SuperScript III platinum One-step qRT-PCR system (Invitrogen, Carlsbad, USA). RT-qPCR assays were performed using conserved primers designed by multiple alignment of available bat DPP4 gene sequences (Table 2), using β-*actin* for normalization. cDNA was amplified in a LightCycler 480 (Roche, Basel, Switzerland) with 25 μl reaction mixtures containing 2× reaction mix, 5 μl RNA, 25 μM ROX reference dye, 50 μM primers and 10 μM probe at 50°C for 30 min, then 95°C for 2 min followed by 50 cycles of denaturation, annealing and extension. Experiments were performed in triplicates, and results were expressed as the mean expression level of DPP4/*β-actin.* The relative expression between different bat cells was then calculated by ΔΔCt method.

### tpDPP4 overexpression in *T. pachypus* cells

*T. pachypus* DPP4 (tpDPP4) sequence was cloned into pLenti7.3/V5-TOPO vector (Invitrogen). The construct was transfected into 293FT cells together with ViraPower Packaging Mix (Invitrogen) using Lipofectamine 2000 (Life Technologies, Carlsbad, USA). Lentivirus was harvested from the supernatant and concentrated using PEG-it (System Biosciences, Palo Alto, USA). *T. pachypus* cells resistant to MERS-CoV were transduced using the concentrated lentivirus for tpDPP4 overexpression and were subsequently subjected to inoculation with 1 MOI of MERS-CoV.

### Nucleotide sequence accession numbers

The nucleotide sequences of bat DPP4 obtained from this study have been deposited in the GenBank sequence database under accession numbers MH345671-MH345676.

## ACKNOWLEDGEMENTS

We thank C. T. Shek, Agriculture, Fisheries and Conservation Department, the Government of the HKSAR, for capture of bats for bat cell development and expert opinion. This work is partly supported by Theme-Based Research Scheme (T11-707/15-R) and Research Grant Council Grant, University Grant Council; Health and Medical Research Fund of the Food and Health Bureau of HKSAR; University Development Fund and Special Research Achievement Award, The University of Hong Kong; Consultancy Service for Enhancing Laboratory Surveillance of Emerging Infectious Disease for the HKSAR Department of Health; The Collaborative Innovation Center for Diagnosis and Treatment of Infectious Diseases, the Ministry of Education of China; National Science and Technology Major Project of China (grant number 2012ZX10004213). Views expressed in this paper are those of the authors only, and may not represent the opinion of the AFCD or the Government of the HKSAR.

